# Temperature-Sensitive Contacts in Disordered Loops Tune Enzyme I Activity

**DOI:** 10.1101/2022.06.18.496683

**Authors:** Daniel Burns, Aayushi Singh, Vincenzo Venditti, Davit A Potoyan

## Abstract

Homologous enzymes with identical folds often exhibit different thermal and kinetic behaviors. Understanding how enzyme sequence encodes catalytic activity at functionally optimal temperatures is a fundamental problem in biophysics. Recently it was shown that the residues that tune catalytic activities of thermophilic/mesophilic variants of the C-terminal domain of bacterial Enzyme I (EIC) are largely localized within disordered loops, offering a model system with which to investigate this phenomenon. In this work, we employ molecular dynamics simulations and mutagenesis experiments to reveal a mechanism of sequence-dependent activity tuning of EIC homologs.

We find that a network of contacts in the catalytic loops is particularly sensitive to changes in temperature, with some contacts exhibiting distinct linear or non-linear temperature-dependent trends. Moreover, these trends define structurally clustered dynamical modes and can distinguish regions that tend toward order or disorder at higher temperatures. Assaying several thermophilic EIC mutants, we show that complementary mesophilic mutations to the most temperature-sensitive positions exhibit the most enhanced activity while mutations to relatively temperature insensitive positions exhibit the least enhanced activities. These results provide a mechanistic explanation of sequence-dependent temperature tuning and offer a computational method for rational enzyme modification.

**Significance:** Temperature affects the catalytic rates of all enzymes. The impact of temperature on the catalytic activity of an enzyme, however, is convoluted from contributions of protein sequence, structure, and dynamics. As such, understanding and designing the molecular features of enzymes which tune catalytic rates at different temperatures remains a fundamental challenge in biophysics. In this work we have employed molecular simulations and mutagenesis experiments to reveal the temperature tuning mechanism of mesophilic and thermophilic homologues of the C domain of bacterial Enzyme l. We find that enzymes can be tuned to their physiological temperatures through a network of temperature-sensitive residue contacts localized in the disordered loops. Furthermore, we find that among temperature-sensitive contacts some exhibit linear and others non-linear dependence on temperature. These clues offer a promising physics-based approach for tuning enzyme activity.

## Introduction

Enzymes are sophisticated catalytic machines, designed by billions of years of natural selection for optimal activity at their physiological temperatures[1–3]. To fold rapidly and function reliably enzyme sequences have evolved to minimize strong energetic frustration, a demand met by forming rigid scaffolds[4,5]. Studies of energy landscapes of proteins however, have shown however, that some degree of frustration is necessary for functional dynamics[6]. Enzyme active sites in particular have been found to be highly frustrated regardless of oligomeric state, topology, or catalytic mechanism[7,8]. Thus, enzymes are shaped to meet conflicting needs for stable folding and functional dynamics which manifests in having both rigid scaffolds and disordered regions. The fact that homologous enzymes can share an identical fold but exhibit drastically different rates of conformational exchange[9–11] suggests an important role for disordered regions in tuning temperature-dependent dynamics. Identifying this temperature tuning mechanism is an important step in learning the key design principles of thermostable enzymes.

A homologous mesophilic-thermophilic enzyme pair is an ideal system for investigating this activity tuning mechanism. To this end, we employed the well characterized Enzyme I C-terminal domain (EIC) (Fig. 1).

**Fig. 1.**
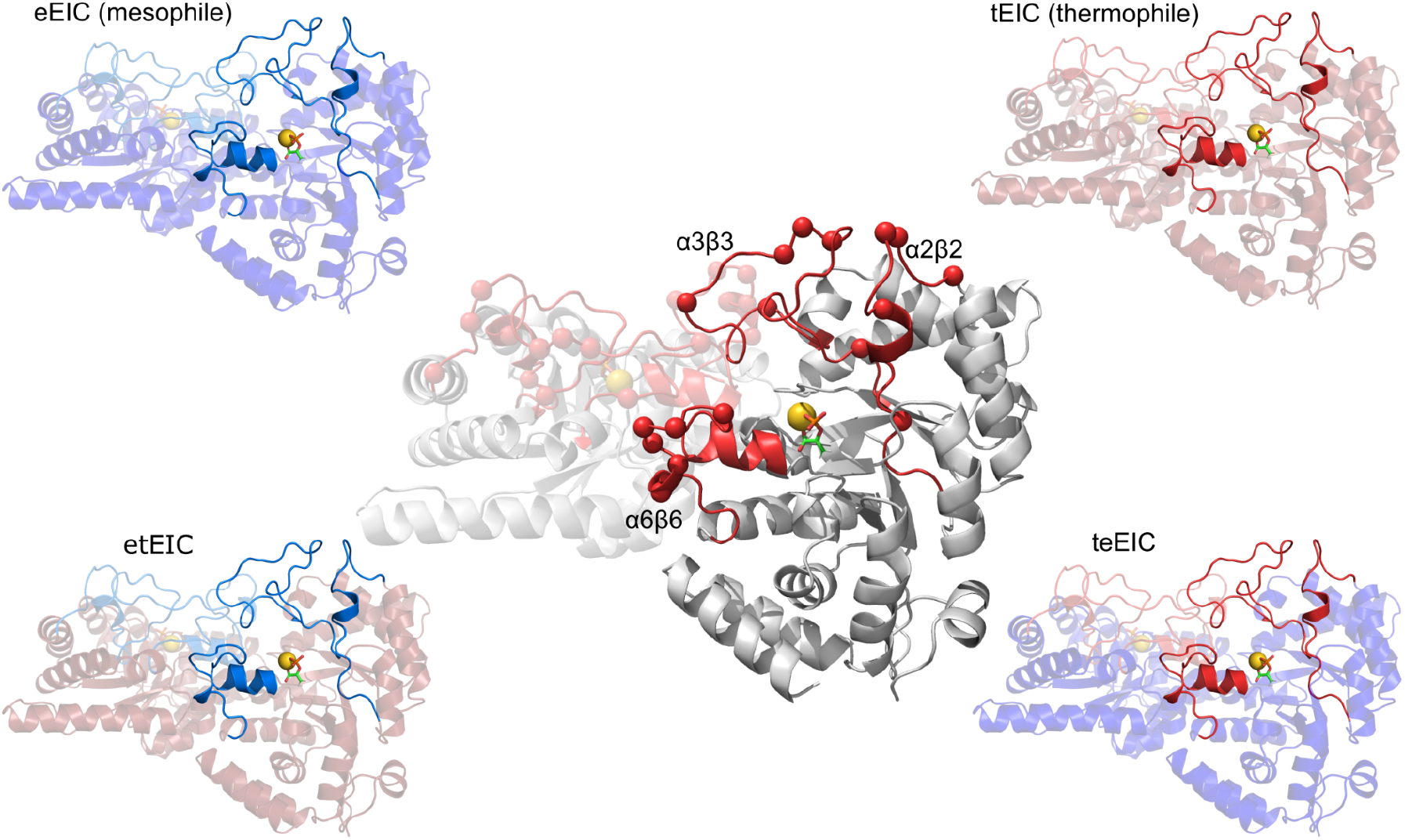
Top: EIC Hybrid construction from wild type homologs. In the center is the EIC structure with 18 of the natural substitutions highlighted as red spheres and names of loops show on top. upper: eEIC; a mesophilic homolog of EIC. upper right:s tEIC; a thermophilic homolog of EIC. bottom left: etEIC; a hybrid formed with a thermophilic scaffold and mesophilic loops.bottom right:teEIC; a hybrid formed with a mesophilic scaffold and thermophilic loops.

Here we used the mesophilic EIC from *Escherichia coli* (eEIC) with a physiological temperature of 37°C (for the natural, full-length enzyme), its structurally-identical thermophilic counterpart from *Thermoanaerobacter tengcongensis* (tEIC) with a physiological temperature of 65°C (full-length), and two hybrids composed of each wild types’ scaffold and disordered catalytic loops swapped with one another (Fig. 1). That is, teEIC contains the thermophile’s loops on the mesophile’s scaffold and etEIC contains the mesophile’s loops on the thermophile’s scaffold. [12]. There are 21 residue substitutions that distinguish the mesophilic and thermophilic loops and importantly, all of the catalytic residues are conserved and there are no deletions or insertions.

The C-terminal dimer catalyzes a phosphoryl transfer reaction from phosphoenolpyruvate (PEP) to water, a reaction that is dependent on the disordered loops assuming a compact conformation[13]. The loops can also assume a catalytically incompentent expanded conformation which involves a large magnitude motion of the α3β3 loop away from the active site. The differences in activity between the wild types can largely be explained by the populations of the expanded or compact conformation determined as a function of the distance between K340 (on the α3β3 loop) and PEP[12].

Previous work demonstrated that the catalytic loops of etEIC exhibit the same expanded to compact conformational exchange rate as the wild type mesophile at 37°C, indicating that this hybrid’s loop dynamics are encoded by the mutations exclusively within the loops [12]. This implies that, in the presence of a stable scaffold, catalytically relevant dynamics can be tuned with a local subset of mutations. This is consistent with other studies demonstrating the amenability of thermophilic enzymes to modification[14]. In addition to shared dynamics, etEIC and eEIC have similar catalytic rates[12,15], implying that these dynamics underlie catalysis.

With the loop dynamics isolated, we can easily detect the effects of individual mutations on activity. We hypothesize that the most temperature-sensitive inter-residue contacts serve to tune the enzyme’s activity to its physiological temperature. To test our hypothesis we sampled catalytic loop conformational ensembles across a wide range of temperatures for each EIC homolog using Hamiltonian Replica Exchange Molecular Dynamics (HREMD) simulations. We then calculated the contact frequencies of all loop residue pairs at each temperature and applied Principal Component Analysis (PCA) to extract the most temperature-sensitive contacts via their loading scores. Finally, we used these most sensitive contact pairs as a guide to design mutant tEIC enzymes based on the natural substitutions in eEIC. Many of these mutants returned activities corresponding well with their PCA based temperature sensitivity rankings, particularly those involving the most temperature sensitive positions. These results strongly support our hypothesis and the utility of our analysis method which we expect to be a valuable research tool for enzyme design.

## Results

### Identifying the temperature-sensitive contacts in enzyme loops

In order to identify the contacts in the tEIC catalytic loop conformational ensemble which are most sensitive to temperature changes, we simulated the enzyme across 20 effective temperatures ranging from 310K° to 510K°. Sampling is done by Hamiltonian Replica Exchange Molecular Dynamics (HREMD), a sampling scheme also known as REST2 (Replica exchange with solute scaling) [16]. The objective of HREMD simulations is to enhance conformational sampling by reducing the strength of the non-bonded interactions involving protein atoms. While the entire system is not heated, the effect of Hamiltonian perturbation is similar and has been shown to reproduce melting curves and folding temperatures with high accuracy[17]. From the sampled MD trajectories we calculated all loop residue contacts with cutoff distances specific to the chemical groups involved [18]. Contact records collected from equilibrium sampling at different temperatures are converted to a contact frequency dataset which describes the percentage of the simulation in which a given residue pair is in contact. The contact frequency datasets have (NT, NP) shape where NT (number of rows) corresponds to temperatures sampled and NP (number of columns) corresponds to contacting residue pairs. An example dataset is available in supplementary figure (SI 1). To reduce the dimensionality of the dataset and to rank the temperature sensitivity of specific residue interactions we employed Principal Component Analysis (PCA).

### Classifying temperature dependence of contacts

Here we used PCA on the contact frequency dataset to identify and rank the residue-residue interactions which are most sensitive to temperature variation. PCA is a linear dimensionality reduction technique that takes in a dataset with N features and returns a set of N orthogonal eigenvectors with corresponding eigenvalues. Eigenvalues of PCs in the present context quantify the degree of temperature-dependent variance of various collective residue contact making/breaking modes [Fig 2AB]. The contact pairs’ loading scores (components of eigenvector) describe the contribution of the initial variables (contact frequencies) to a PC’s explained variance (Fig. 2C). In other words, loading scores provide a natural ranking system for a contact’s temperature sensitivity in terms of the dominant trend described by a given PC.

**Fig. 2.**
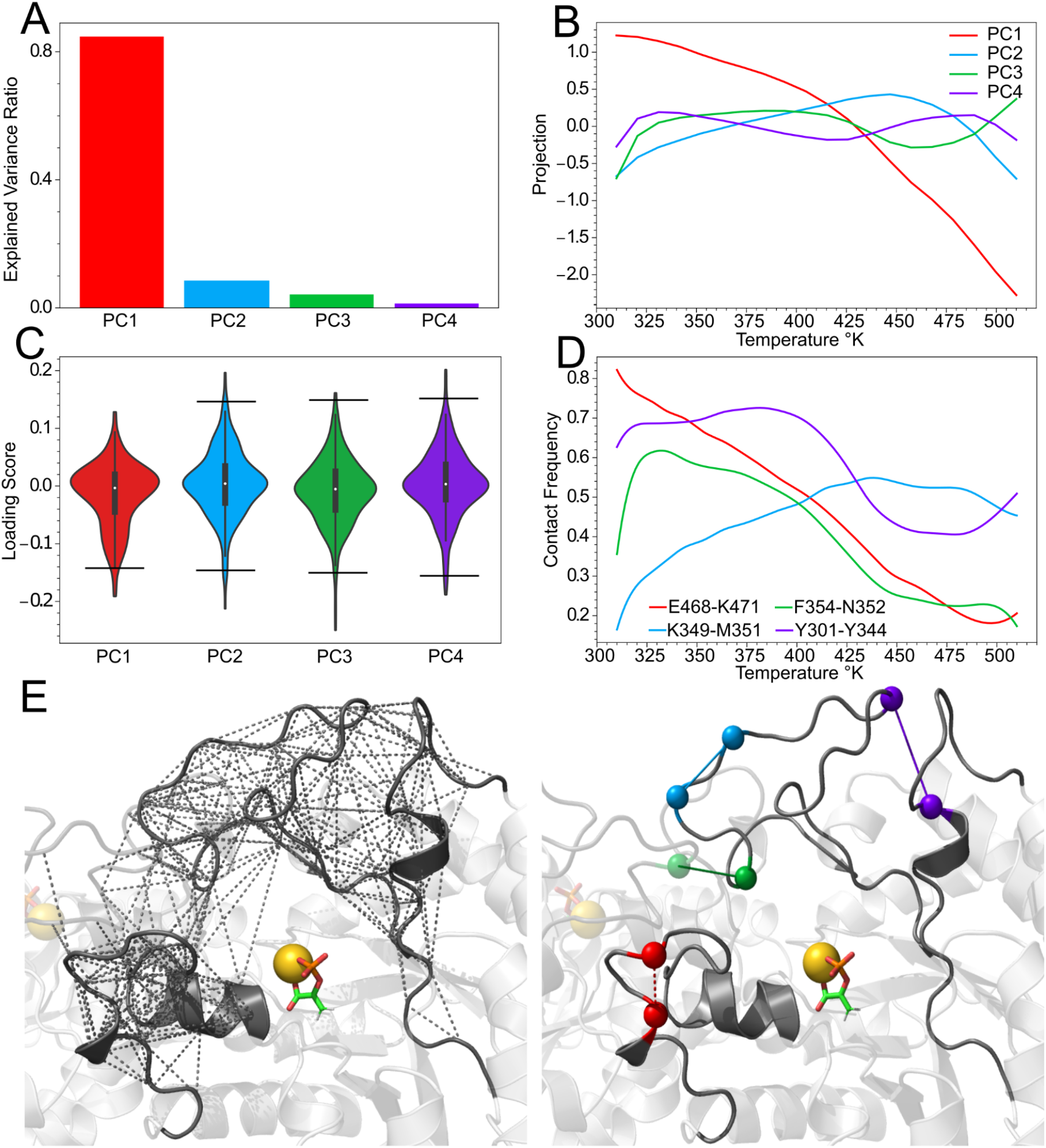
(A) tEIC contact frequency explained variance by PC. (B) PC projection vs effective loop temperature. (C) Histograms of PC loading scores with bars indicating the cutoff for the 98th percentile of loading score absolute values. (D)Contact frequency vs temperature for a selection of highly temperature-sensitive contacts from PCs 1-4.(E) Left: All the averaged contacts depicted on one subunit before PCA categorization and ranking. Right: The most temperature-sensitive contacts distilled from the left image.

Our hypothesis predicts that contacts with very high sensitivity to temperature have the most influence on activity. Here we identified a highly temperature-sensitive contact as occurring in the 98th percentile of the normalized loading score distributions. These include the top 7 contacts on each PC (Fig S3), falling entirely within the tail end of the loading score histogram (Fig. 2C). Because our previous work demonstrated that the loop dynamics of tEIC are essentially uninfluenced by scaffold interactions (etEIC kex/ kcat) we directed our mutation experiments and focused our simulation analysis on tEIC. While the homologs differ in their individual contact behavior, we noted that they displayed similar PC trends indicating that tEIC’s data is representative of the general behavior of the loops regardless of sequence composition (Fig. S4).

For tEIC, PC 1 describes nearly 85% of the variance and captures a linear change in contact frequencies with temperatures (Fig. 2A-B). This represents the dominant effect of heating, increasing configurational entropy via progressive contact breaking toward denaturation. The next PCs identify the less obvious effects of heating. Since their eigenvalues decay exponentially, we focused on the first four PCs which accounted for over 98% of variance and disregarded the remaining PCs. The first 4 principal component projections are shown in figure 2B and the plots of a selection of highly temperature-sensitive contacts’ frequencies vs temperature from each PC (which reflect the trends shown in 2B) is shown in figure 2D.

PC 2 describes less than 10% of the variation in the data, however it captures an important trend of increasing contact frequencies (Fig. 2B-D), becoming progressively more stable across a wide range of temperatures. PC 3 captures a sharply increasing contact frequency within the physiological temperature range before rapidly decaying at higher temperatures. For tEIC this decay begins around its optimal physiological temperature. PC 4’s trend is similar to that of PC 3 (Fig. 2B) but its high ranking contacts include more naturally substituted residues than PC 3 and was therefore more informative for our experimental purposes (‘Mutation Experiments’ Section). The remaining PC trends are less obvious and begin to resemble noise after the 4th PC.

By applying PCA and loading score ‘filtering’ to the contact frequency dataset, we reduced the initially intractable web of contacts to only the contacts with the strongest response to changes in temperature (Fig. 2E).

### Mapping temperature-sensitive contact networks

A global view of the disordered loop contacts and their PC ranked temperature sensitivities can be visualized with contact maps (Fig 3). On PC1 the highly temperature-sensitive contacts form an obvious cluster on the α6β6 loop, which is also apparent when viewed on the structure (Fig. 3). These high ranked PC1 contacts have a strong tendency to decrease across the full temperature and frequency range.

**Fig. 3.**
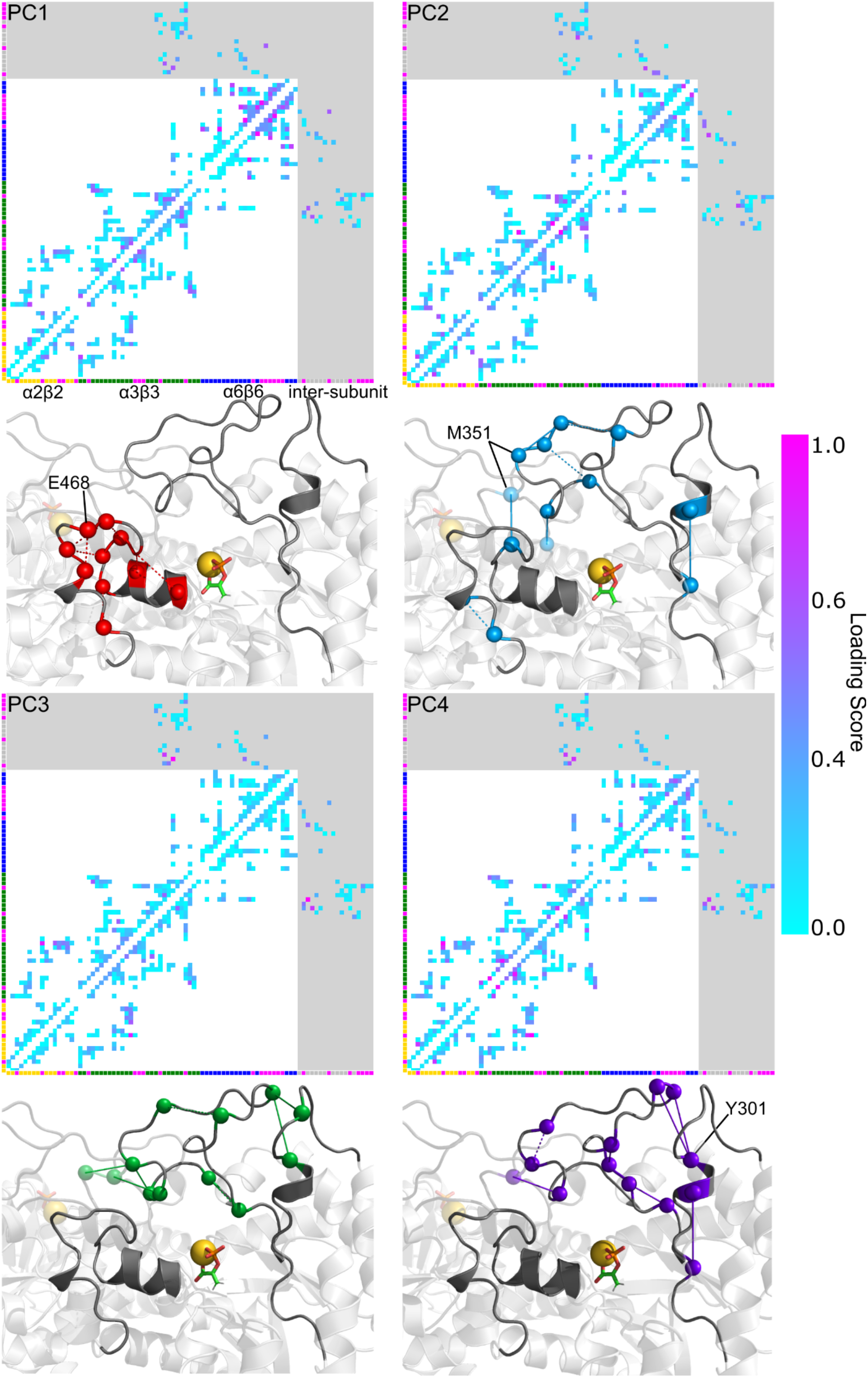
Contact maps showing normalized loading scores (temperature sensitivity) according to the colorbar (right). Corresponding structures are shown with the top 7 most temperature-sensitive contacts for the given PC. Solid lines indicate contacts that have a tendency to increase in frequency in the physiological temperature range. Dashed lines tend to decrease in frequency in the same range. The tick marks on the axes of the contact maps indicate residues and are colored as follows: yellow: α2β2, green: α3β3, blue: α6β6 and magenta indicates a naturally substituted position.

PC2 is the first PC to capture a number of strongly associating contacts with non-linear profiles. On PC2, the highest ranking contact involves M351. This natural substitution is in the middle of 9 conserved residues on the α3β3 loop, particularly isolated relative to other substitutions, and directly involved in the structurally and functionally vital dimer interface[19]. A glutamate in the mesophile, this residue also represents the largest change in hydrophobicity among the loop substitutions. The top sensitive contacts on PCs 3 and 4 are entirely devoid of residues in the α6β6 loop and share similar regions of the structure and display similar contact frequency trends. PC 4’s contacts, however, are shifted more towards the α2β2 loop indicating that it better defines this region than PC 3.

The loading score values decay differently on different PCs resulting in some contacts with comparatively low loading scores being considered as highly temperature-sensitive. For instance, considering the top 7 contacts on each PC results in a high loading score cutoff of 0.8 for PC 4 but a much lower value of 0.6 for PC3. The loading score contact maps offer additional insight as to which contacts are most important within each PC. Additionally, ome contacts can score highly on multiple PCs as shown in the score plots (Fig. S6 “F354-N352” of PC3 and PC4 for instance), making some only marginally better defined by one PC than the other. Nevertheless, having identified the temperature-sensitive contacts within the disordered loops of tEIC, we sought experimental evidence to tell us whether or not they have physical significance.

### Mutation Experiments

The EI homologs provide a known set of function-conserving mutations between the mesophile and the thermophile. If the residues that occur in highly temperature-sensitive contacts are among the natural substitutions, it suggests that these positions have been selected based on their simultaneous tuning of loop dynamics and preservation of catalytic conformational space.

To investigate this, we made point mutations to several of tEIC’s naturally substituted positions with the complementary residues of eEIC. We assayed these mutants for activity by measuring the rate of PEP hydrolysis at 40° C (a temperature at which tEIC is nearly inactive) and comparing their initial rates with etEIC, which provided the hypothetical maximum rate achievable from a subset of mutations[12]. In our model, the hydrolysis rate of PEP indicates how effectively the mutation has enhanced the disordered loop dynamics underlying catalysis. For the natural substitutions that do not appear in the most temperature-sensitive contacts (the top 98th percentile of loading scores on the first four PCs), we expected their complementary point mutation to yield the least enhanced activities.

The assay results ultimately revealed a noteworthy correspondence between several of tEIC’s mutant activities and highest contact temperature sensitivity rankings (Fig. 4B).

**Fig. 4.**
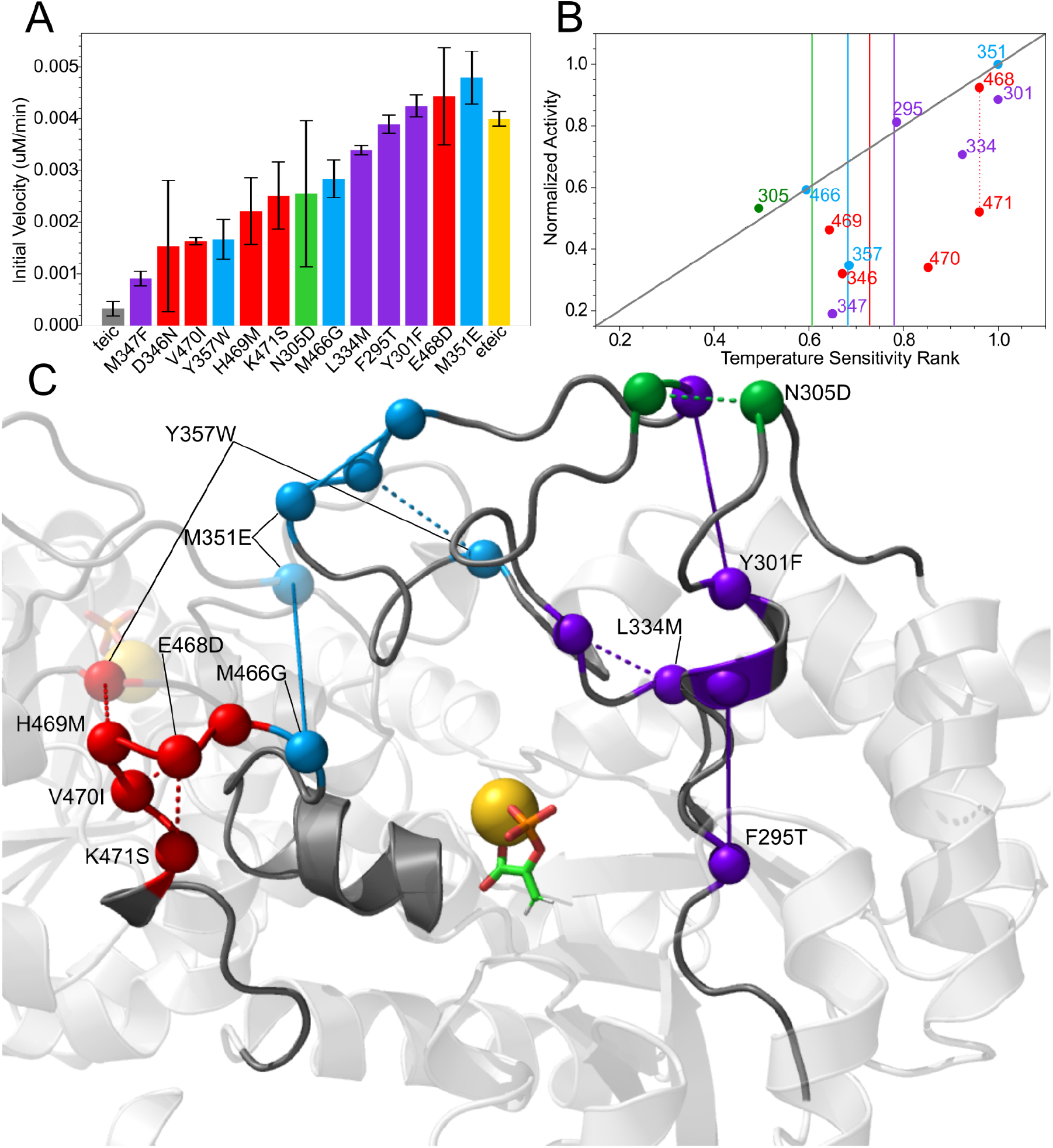
(A) Assay results for tEIC mutations colored according to the PC in which they have the highest loading score contact (red: PC1, blue: PC2, green: PC3, purple: PC4). Colored solid vertical lines show the loading score value cutoff for the 98th percentile for the corresponding PC. (B) Mutant activity vs temperature sensitivity rank on their highest tEIC loading scoring contact PC. dotted line indicates that the loading score for 468 and 471 comes from the same contact (C) Contacts involving residues from the mutant assays colored according to the PC on which their highest loading score occurs (lowest activity enhancing mutations on 346 and 347 contacts not shown).

Remarkably, the mutations that returned the highest activities are indeed participants in the highest ranked substituted contacts and thus, the most temperature-sensitive. The Pearson correlation of the activities and temperature-sensitivities of all the assayed mutants is 0.632.

The top PC1 guided mutants (V470I, K471S, E468D) resulted in mixed activity enhancement (Fig. 4A) with the most enhancing mutation from the PC1 rankings being E468D, guided by the top-ranked substituted contact (2nd overall) involving residue K471.

M351, involved in the highest ranked temperature-sensitive tEIC contact on PC 2, when mutated, resulted in the highest activity enhancement among all of our mutants. Its contact with the conserved K349 exhibits a strong increasing frequency trend, remaining stable well above physiologically relevant temperatures. While having a large impact on activity, very little thermal-stability was sacrificed by this mutation or indeed any of our point mutations relative to the destabilization etEIC exhibits from the full complement of mutations (Fig. S7).

The third highest mutant activity was returned by Y301F whose contact with the conserved Y344 ranks 2nd and 1st on PC 3 and 4 respectively.

The F295T mutant returned the 4th highest activity with its contact to the conserved F299 ranking 4th on PC 2 and in the top 6 of PC 4.

The fifth most active mutant, L334M, ranks 3rd on PC4 with its contact to the conserved I336. The only naturally substituted residue outranking it on this PC was Y301. Residues occurring to the left of their respective cutoff line in figure B were mutated and assayed to compare their rank-activity relationship with the highly-temperature sensitive contact residues. A table of each PC’s top 7 temperature-sensitive contacts indicating which residues are naturally substituted is available in figure S8.

Depicting the experimentally mutated residue’s highest ranking contacts such that they are colored according to the PC on which they have the highest loading scores results in a clear structural clustering [fig 4C] of the different modes defined by the PCs. In terms of naturally substituted residues, the loops are largely characterized by seperate PC modes. Visualizing the contacts according this same color scheme for all of the top contacts (regardless of whether they involve substituted residues or not) on PCs 1-4 reveals that PC3, with primarily conserved residue contacts, best describes the interaction at the interface of the two subunits’ α3β3 loops (SI 9).

## Discussion

Energy landscape theories of proteins have shown that frustration and disorder are evolutionarily conserved features which are necessary for function [7,8]. The nature of interactions which balance the competing needs for having ordered catalytic sites and dynamical disorder, however, is poorly understood. Here we have employed molecular dynamics simulations and in vitro assay experiments on the C domain of bacterial Enzyme I (EIC) to reveal the mechanism underlying activity-tuning in mesophilic (tEIC) and thermophilic enzyme homologues (eEIC).

We find that functionally optimal temperature for a tEIC, critical residue contacts are not accessible in disordered loop motions. As temperature increases, certain contacts break (e.g. PC1), allowing other critical contacts to form (e.g. PC2). Eventually a temperature is reached where the optimal interactions are occurring to cycle the enzyme through catalytically relevant conformations at a rate tuned to the functional needs of the organism. This enhancement can involve a critical contact whose altered dynamics promote relevant conformational sampling. By mutating a specific residue involved in the critical contact the enzyme can sample the catalytic conformations at a lower temperature.

Furthermore, we present a computational method for ranking temperature-sensitive contacts and identifying residues critical for temperature-sensitive behavior of enzymes. We find highly temperature-sensitive contacts displaying linear and non-linear temperature dependent frequency trends. These trends define structurally clustered dynamical modes and can distinguish contacts which tend toward order or disorder at higher temperatures. Assaying several thermophilic EIC mutants, we show that complementary mesophilic mutations to the most temperature-sensitive positions, identified by their high loading scores, exhibit the most enhanced activity while mutations to relatively temperature insensitive positions exhibit the least enhanced activities.

In contrast to loading scores, we find that a larger eigenvalue is not more predictive of activity enhancing mutations (Fig. 2B) as exemplified by assay results of M351E (PC2) and T301F (PC4) (Fig. 4B). This is because eigenvalues are a measure of collective contact dynamics. Thus PC1 captures the collective melting trend which is the best representation of the entire protein’s response to elevated temperatures and thus while eigenvalue modes describe subtler trends perhaps more specific to catalytic function.

Beyond the activity tuning application outlined here, a number of applications for temperature-sensitive contact analysis are possible. We speculate that regions involving temperature-sensitive contacts might serve as drug targets. Since these residues lie at the center of a dynamical mode and are important for activity, it follows that their activity could be inhibited by disrupting their dynamics with the binding of small molecules. The contact analysis presented here could also be extended to identify pH sensitive contacts by way of variable protonation state simulation schemes or analyzing multiple independent simulations involving different residue tautomers. Finally, we envision temperature-sensitive contact analysis providing insights into mechanisms of systems such as ion channels, biomolecular condensates, and functionally relevant dynamics of biomolecular assemblies in general.

## Materials and Methods

### Simulation Protocol

Crystal structure for tEIC was obtained from PDB entries 2BG5 (tEIC). Molecular dynamics simulations were carried out using Gromacs 2018.8 (The last 50ns of teEIC was done with 2019 on Expanse) utilizing the leapfrog integration method with a 2fs timesteps [20–23]. Short range electrostatic and Lennard-Jones interactions were calculated with a plain coulomb cutoff of 1.2 nm. The Particle mesh Ewald electrostatics scheme was employed for long range electrostatics with a grid spacing of 0.16 nm [24]. Bonds to hydrogen were constrained using the LINCS algorithm [25]. The system was solvated in a dodecahedral simulation box with TIP3P water molecules and neutralized with 100mM NaCl. The solvent and solute were separately coupled to a modified Berendsen thermostat (Velocity Rescale) with a reference temperature of 310K. Simulations included PEP to simulate the dynamics relevant to the rate limiting chemical step. The system was described according to the CHARMM 36 force field with the PEP ligand molecule parameters provided by cgenff [26–28]. The system was energy minimized with the steepest descent minimization and equilibrated in the NVT and NPT ensembles for 100 ps.

### Hamiltonian Replica Exchange Molecular Dynamics Simulations

To efficiently sample the temperature dependent conformational ensembles of Enzyme I disordered loops we employed Hamiltonian Replica Exchange Molecular Dynamics (HREMD) [29–31], specifically the REST2 solute tempering scheme. The choice of the Hamiltonian method was motivated by the size of the system which makes application of temperature replica exchange (TREMD) computationally costly. The residues 291-309, 332-360, and 454-477 within the active site loops of the constructs were selected for Hamiltonian perturbation which includes 18 of the 21 substituted positions in the hybrid enzyme (the remaining three being on short linkers not enhanced with the Hamiltonian treatment). Residues within these loops were identified as central to catalysis in previous work via NMR and structure visualization[12]. A total of 20 replicas with an effective temperature range of 310-510° K was chosen as it gave an exchange rate around 25%. Hamiltonian replica exchange simulations were performed with gromacs 2018.8 patched with Plumed 2.5.5 on Xsede’s Comet and Expanse HPC systems [32,33][34]. Exchanges were attempted every 800 timesteps. Trajectory snapshots were extracted every 10ps. Simulations were run for 200ns which generated an effective 400ns sampling by combining contact data from the two subunits. A wall potential restricting the PEP molecule to within 6.5Å of cysteine 502, a residue centered beneath the binding site, was employed to prevent unbinding of PEP in the high temperature replicas and ensure enhanced sampling of the relevant motions associated with the bound substrate. A weak 50 kJ/mol dihedral restraint was also applied to the CCOP dihedral angle of PEP to bias the hypothetically catalytically competent PEP conformation.

### Contact analysis

We calculated the inter-atomic contacts and contact frequencies of the loop residues using GetContacts[35]. Contacts between directly adjacent residues were discarded as well as contacts whose frequencies did not exceed 5% across the 20 replicas. The frequencies for both subunits were averaged and contacts not occurring in both subunits were also discarded. This left approximately 300 contacts for analysis. All subsequent calculations were performed on the averaged values. The PCA loading scores were normalized according to (absolute value/maximum absolute value on the PC). The 98th percentile of these scores was calculated to determine which contacts to consider highly temperature-sensitive. Molecular visualizations were produced with pymol.

### Mutations

Point mutations were performed according to the Agilent Quickchange II protocol.

### Enzyme Expression

Plasmids for etEIC and tEIC constructs with His and EIN solubility tags were transformed into BL21 cells and cultured on LB media overnight. All culture media contained 100μg/ml ampicillin. Single colonies were grown in 20ml of LB media overnight at 37°C and used to inoculate 1L LB cultures. These were grown to an OD600 of 0.6 before cooling to 20°C and adding 0.238g of IPTG for overnight expression. Cultures were spun at 4,000×g for 30 minutes and resultant pellets frozen at −80°C.

### Enzyme Purification

Pellets were thawed on ice and cells lysed and homogenized at (1200 psi) via Avestin Emulsiflex in 20ml of buffer containing 50mM Tris pH 8, 300mM NaCL, and 5mM imidazole. Lysate was centrifuged at 37,000 × g for 60 minutes followed by filtration through a 10kDa membrane and loaded on to a Bio-Rad Nuvia Ni-charged column. Protein was eluted with a 500mM Imidazole buffer applied as a gradient. Eluted protein was buffer-exchanged with the original buffer and incubated with TEV protease to remove the solubility tag. Samples were filtered and again passed through the Ni column where the solubility tag was retained and the eluted enzyme collected. The fractions were buffer exchanged with 20mM Tris pH 8, 1mM EDTA, and 2mM DTT before being applied to a Bio-Rad EnrichQ column and eluted with a 1M NaCL buffer gradient. The purified product was buffer exchanged with water and stored at −80°C.

### Activity assays

(A280/ uvVis) values were used with molar extinction coefficients obtained from (expasy protparam) sequence analysis to determine protein concentrations. Enzymes were diluted to a concentration of 50 uM in an NMR buffer containing 3mM PEP and 1mM TSP. Reactions at 40°C were monitored by the decrease in PEP (vinyl hydrogen?) peak intensity at 5 minute intervals on a 700MHz Bruker NMR over the course of 2 ½ hours. PEP concentration was normalized against TSP peak intensity. Enzyme concentrations from the experimental samples were normalized against the etEIC experimental sample concentration determined by SDS PAGE band intensities with imageJ [36]. The average concentration (band intensities) from the two etEIC replicates was divided by the mutant’s band intensity to produce a normalization coefficient for each corresponding rate.

## Data availability

All study data are included in the article and/or SI. The scripts used for analyzing molecular dynamics trajectories and for creating all the figures reported in this study are available at https://github.com/PotoyanGroup/Temp-sens-contacts

## Acknowledgements

This work was supported by funds from the National Institute of General Medical Sciences with grant no. R35GM133488 [to V. V.] and grant no. R35 GM138243 [to D.P.]

