## Supplemental Information for "Temperature-Sensitive Contacts in Disordered Loops Tune Enzyme I Activity"

### Supplementary Information

| | A:LYS:471-A:TYR:474 | A:HIS:469-B:TYR:357 | A:ARG:304-A:MET:302 | A:LEU:355-B:PHE:354 | A:LEU:342-A:TYR:301 | A:GLN:458-A:PHE:354 | A:LEU:355-B:ALA:462 | A:GLU:298-A:MET:302 | A:LEU:355-A:PRO:353 |
|---|---|---|---|---|---|---|---|---|---|
| 310 | 0.8015 | 0.6485 | 0.9705 | 0.3790 | 0.5995 | 0.6335 | 0.4150 | 0.9180 | 0.9670 |
| 318 | 0.7895 | 0.6085 | 0.9690 | 0.4170 | 0.5955 | 0.5535 | 0.4435 | 0.9015 | 0.9305 |
| 327 | 0.7800 | 0.5780 | 0.9680 | 0.4270 | 0.5870 | 0.5185 | 0.4760 | 0.8935 | 0.9240 |
| 335 | 0.7655 | 0.5600 | 0.9680 | 0.4225 | 0.5770 | 0.5020 | 0.4855 | 0.8845 | 0.9225 |
| 344 | 0.7460 | 0.5425 | 0.9690 | 0.4115 | 0.5710 | 0.4820 | 0.4980 | 0.8785 | 0.9220 |
| 353 | 0.7290 | 0.5255 | 0.9685 | 0.4085 | 0.5710 | 0.4720 | 0.4985 | 0.8740 | 0.9250 |
| 363 | 0.7160 | 0.5065 | 0.9690 | 0.4085 | 0.5700 | 0.4690 | 0.4955 | 0.8655 | 0.9280 |
| 372 | 0.7090 | 0.4940 | 0.9690 | 0.3985 | 0.5640 | 0.4685 | 0.5050 | 0.8580 | 0.9315 |
| 382 | 0.6980 | 0.4845 | 0.9685 | 0.3915 | 0.5590 | 0.4585 | 0.5145 | 0.8470 | 0.9265 |
| 392 | 0.6845 | 0.4775 | 0.9675 | 0.3725 | 0.5455 | 0.4515 | 0.5260 | 0.8365 | 0.9185 |
| 403 | 0.6755 | 0.4655 | 0.9670 | 0.3590 | 0.5210 | 0.4315 | 0.5490 | 0.8230 | 0.9070 |
| 414 | 0.6525 | 0.4490 | 0.9655 | 0.3410 | 0.5010 | 0.4060 | 0.5650 | 0.8065 | 0.8935 |
| 425 | 0.6165 | 0.4230 | 0.9650 | 0.2975 | 0.4910 | 0.3755 | 0.5765 | 0.7835 | 0.8760 |
| 436 | 0.5590 | 0.3990 | 0.9640 | 0.2655 | 0.4885 | 0.3410 | 0.5800 | 0.7590 | 0.8565 |
| 447 | 0.4915 | 0.3670 | 0.9635 | 0.2395 | 0.4920 | 0.3285 | 0.5660 | 0.7370 | 0.8470 |
| 459 | 0.4380 | 0.3445 | 0.9660 | 0.2315 | 0.4895 | 0.3215 | 0.5430 | 0.7120 | 0.8420 |
| 471 | 0.4090 | 0.3020 | 0.9660 | 0.2200 | 0.4705 | 0.3155 | 0.5190 | 0.6750 | 0.8290 |
| 484 | 0.4065 | 0.2300 | 0.9675 | 0.2215 | 0.4485 | 0.3060 | 0.4765 | 0.6285 | 0.8050 |
| 497 | 0.4510 | 0.1765 | 0.9670 | 0.2195 | 0.4075 | 0.2995 | 0.4335 | 0.5470 | 0.7845 |
| 510 | 0.4685 | 0.1430 | 0.9685 | 0.1855 | 0.3670 | 0.2795 | 0.3630 | 0.5010 | 0.7850 |

**Fig. S1**. Example contact frequency data. Contact pairs IDs are labeled above each column in the format of: ‘Chain ID 1: Residue Name 1: Residue Index 1 - Chain ID 2: Residue Name 2: Residue Index 2’. Different chain IDs indicate that the contact is occurring between the two subunits. Columns contain contact frequencies of a contact pair with each row giving a frequency at a different temperature. Row indices are temperatures in K° based on the plumed_scaled_topologies effective temperature calculation. Values are averaged from both subunits.

**X**: active site **X**: conserved residues —EIN —EIC - - - active site loops *: mutations

```
eEI   1    MISGILASPGIAFGKALLLKEDEIVIDRKKISADQVDQEVERFLSGRAKASAQLETIKTK   60
tEI   1    MLKGVAASPGIAIGKAFLYTKEKVTINVEKIEESKVEEEIAKFRKALEVTQEEIEKIKEK   60

eEI   61   AGETFGEEKEAIFEGHIMLLEDEELEQEIIALIKDKHMTADAAAHEVIEGQASALEELDD   120
tEI   61   ALKEFGKEKAEIFEAHLMLASDPELIEGVENMIKTELVTADNAVNKVIEQNASVMESLND   120

eEI   121  EYLKERAADVRDIGKRLLRNILGLKIIDLSAIQDEVILVAADLTPSETAQLNLKKVLGFI   180
tEI   121  EYLKERAVDLRDVGNRIIENLLGVKSVNLSDLEEEVVVIARDLTPSDTATMKKEMVLGFA   180

eEI   181  TDAGGRTSHTSIMARSLELPAIVGTGSVTSQVKNDDYLILDAVNNQVYVNPTNEVIDKMR   240
tEI   181  TDVGGRTSHTAIMARSLEIPAVVGLGNVTSQVKAGDLVIVDGLEGIVIVNPDEKTVEDYK   240

eEI   241  AVQEQVASEKAELAKLKDLPAITLDGHQVEVCANIGTVRDVEGAERNGAEGVGLYRTEFL   300
tEI   241  SKKESYEKKVEGLKQLKDLPAETPDGKKVMLAANIGTPKDVASALANGAEGVGLFRTEFL   300
                                                                 β2α2 loop
eEI   301  FMDRDALPTEEEQFAAYKAVAEACGSQAVIVRTMDIGGDKELPYMNFPKEENPFLGWRAI   360
tEI   301  YMDRNSLPSEEEQFEAYKEVVEKMGGRPVTIRTLDIGGDKELPYLDMPKEMNPFLGYRAI   360
           (res. 296-309)                          β3α3 loop (res. 332-360)
eEI   361  RIAMDRKEILRDQLRAILRASAFGKLRIMFPMIISVEEVRALRKEIEIYKQELRDEGKAF   420
tEI   361  RLCLDRPDIFKTQLRAILRASAYGNVQIMYPMISSVEEVRKANSILEEVKAELDREGVKY   420

eEI   421  DESIEIGVMVETPAAATIARHLAKEVDFFSIGTNDLTQYTLAVDRGNDMISHLYQPMSPS   480
tEI   421  DKEIKVGIMVEIPSAAVTADILAKEVDFFSIGTNDLTQYTLAVDRMNEHVKEYYQPFHPA   480
                                                    β6α6 loop (res. 454-477)
eEI   481  VLNLIKQVIDASHAEGKWTGMCGELAGDERATLLLLGMGLDEFSMSAISIPRIKKIIRNT   540
tEI   481  ILRLVKMVIDAAHKEGKFAAMCGEMAGDPLAAVILLGLGLDEFSMSATSIPEIKNIIRNV   540

eEI   541  NFEDAKVLAEQALAQPTTDELMTLVNKFIEEKTIC   575
tEI   541  EYEKAKEIAEKALNMSEAREIEKMMKDVI--KDIG   573
```

**Fig. S2.** Sequence alignment of full length eEI and tEI showing that the loops (dotted underline) contain identical catalytic residues and number of residues defining the loops. Black letters indicate natural substitutions. The EI C domain is underlined in red.

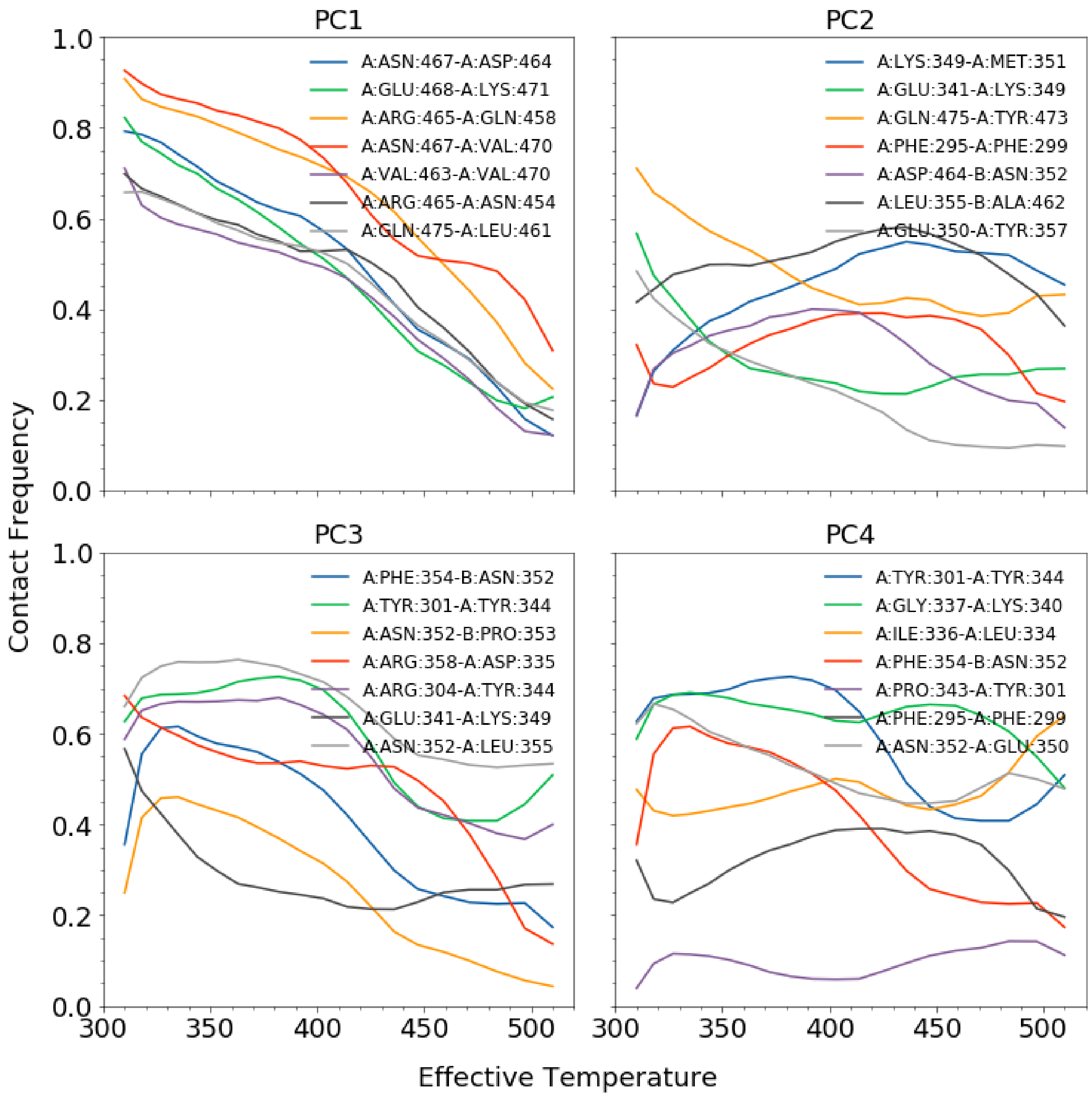


**Fig. S3**. Top 7 most temperature sensitive contact frequencies vs temperature on the first 4 PCs.

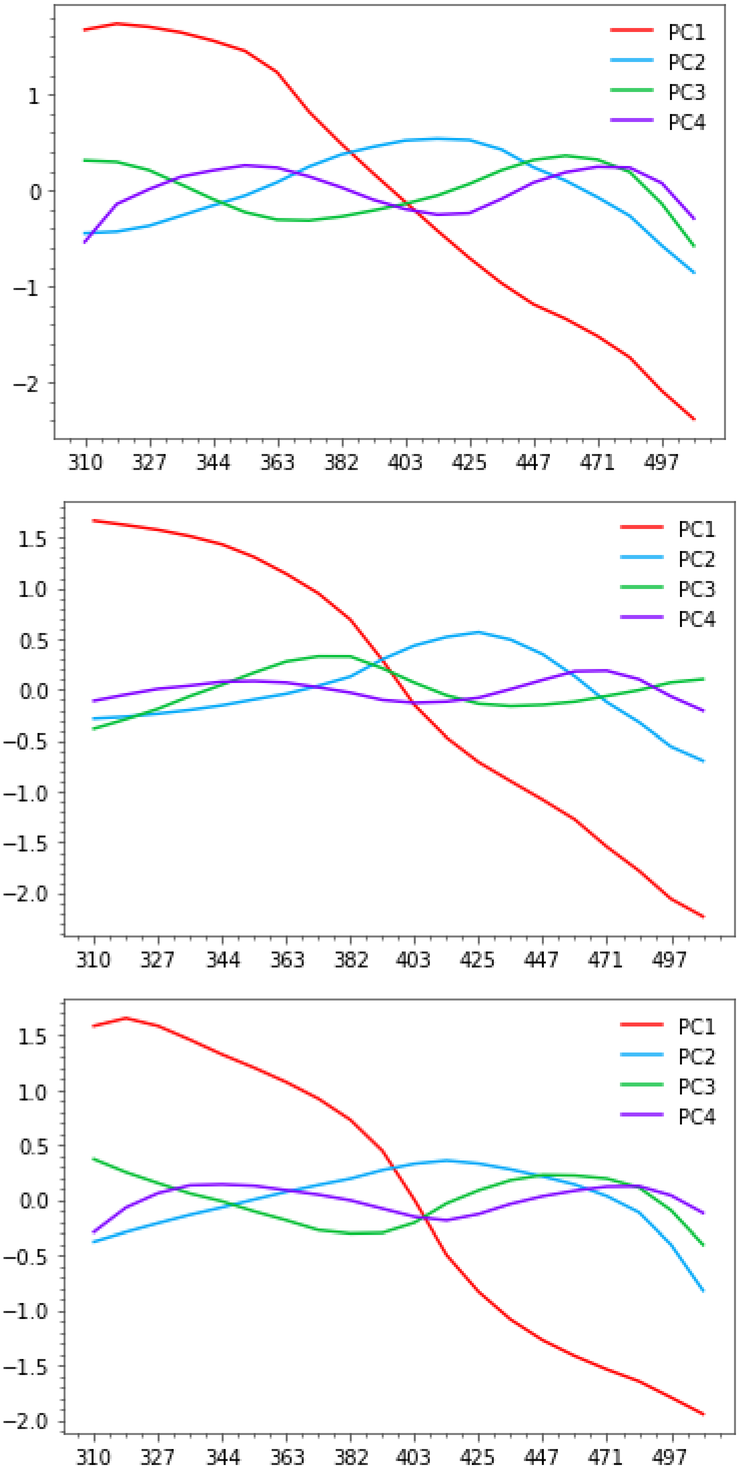


**Fig. S4.** Plots of homolog contact frequency PC projections. Top to bottom: eEIC, eTEIC, teEIC. All systems were run at the same temperature ranges so subtle differences in trends are expected as all of these enzymes show higher activity at lower temperatures than tEIC.

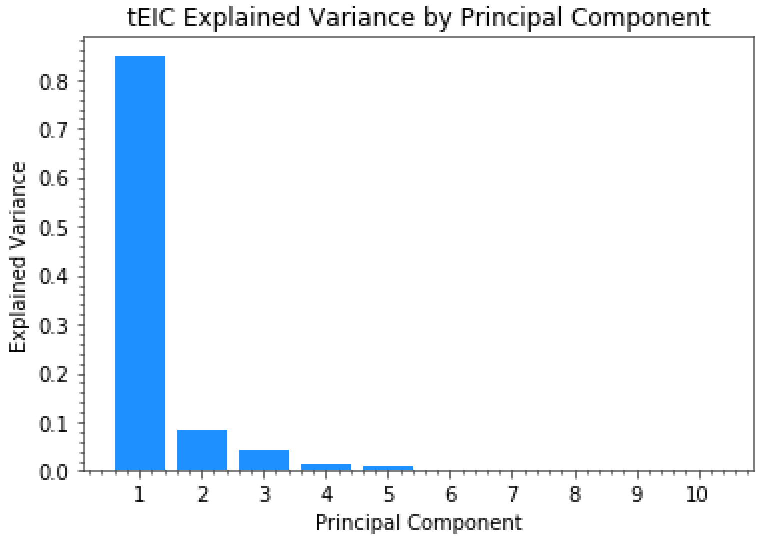


**Fig. S5.** Explained variance of tEIC enzyme ordered by PC eigenvalue magnitude.

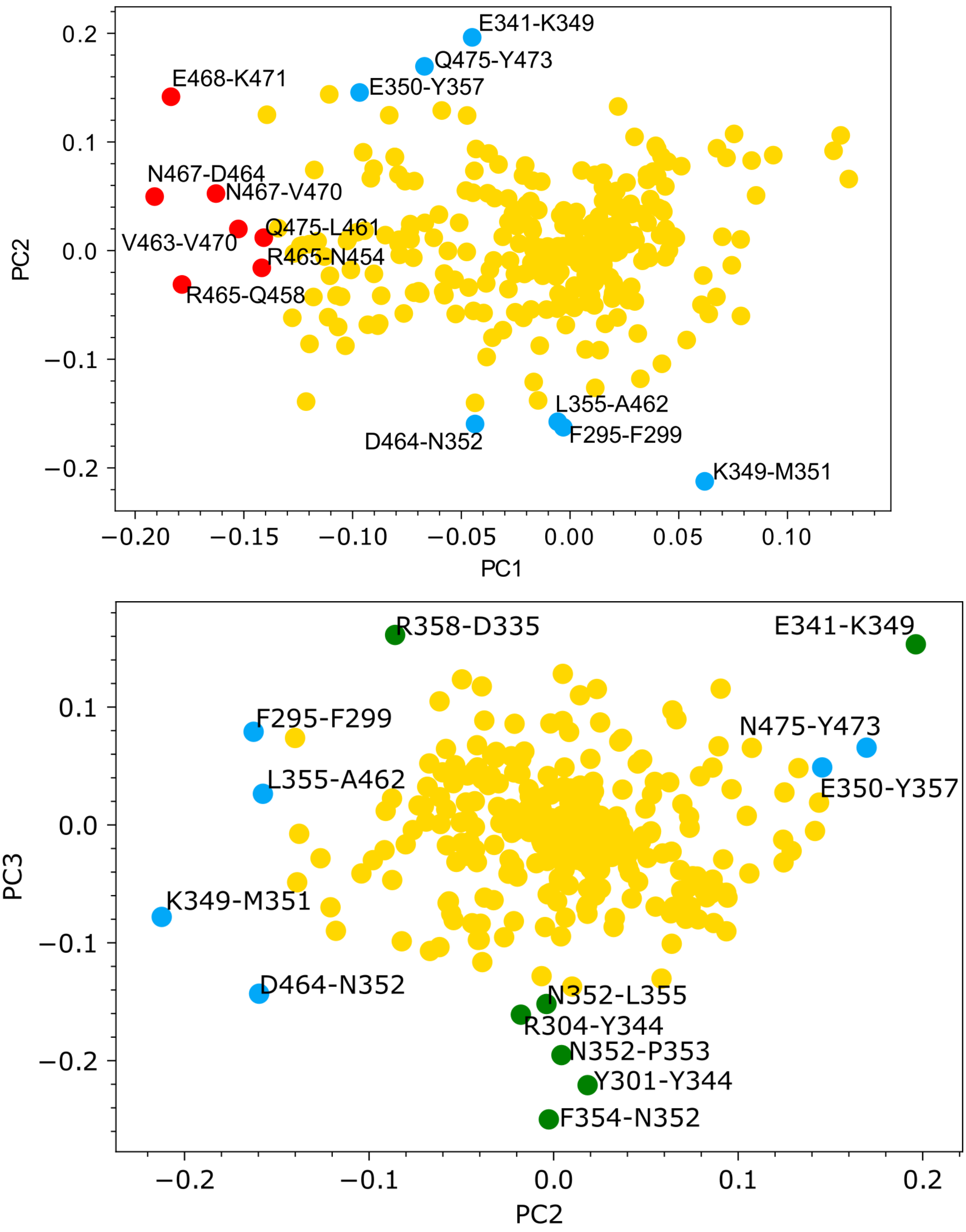
E341-K349
Q475-Y473
E350-Y357
E468-K471
N467-D464
N467-V470
Q475-L461
V463-V470
R465-N454
R465-Q458
L355-A462
F295-F299
D464-N352
K349-M351
PC1
PC2
R358-D335
E341-K349
F295-F299
N475-Y473
L355-A462
E350-Y357
K349-M351
D464-N352
N352-L355
R304-Y344
N352-P353
Y301-Y344
F354-N352
PC2
PC3

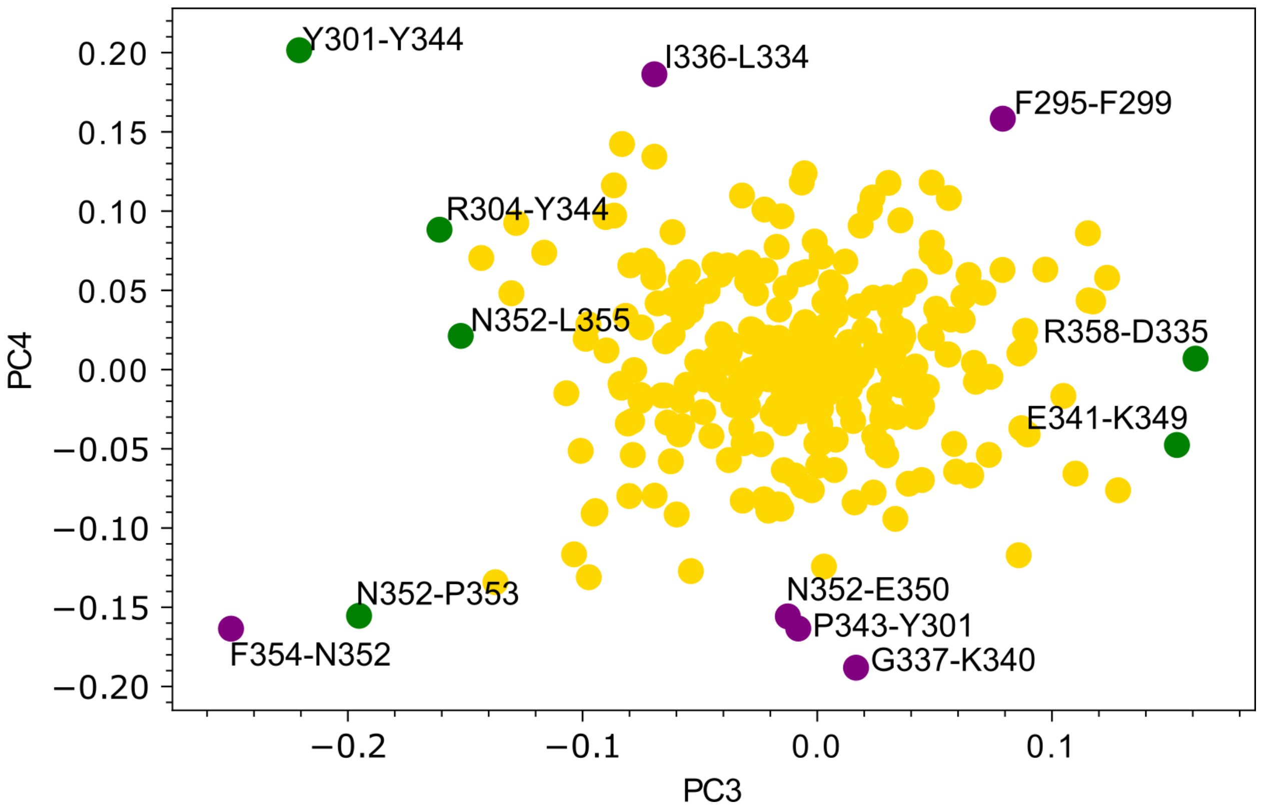


**Fig. S6.** Top 7 temperature-sensitive contacts from the two PCs indicated on the axes plotted according to their (non-normalized) loading scores on each PC. (Red: PC1, blue: PC2, green: PC3, purple: PC4). These highlight that the highly sensitive contacts occur on the extremes of the mode identified by the PC. Points near the corners are considered highly sensitive according to the trend described by both modes.

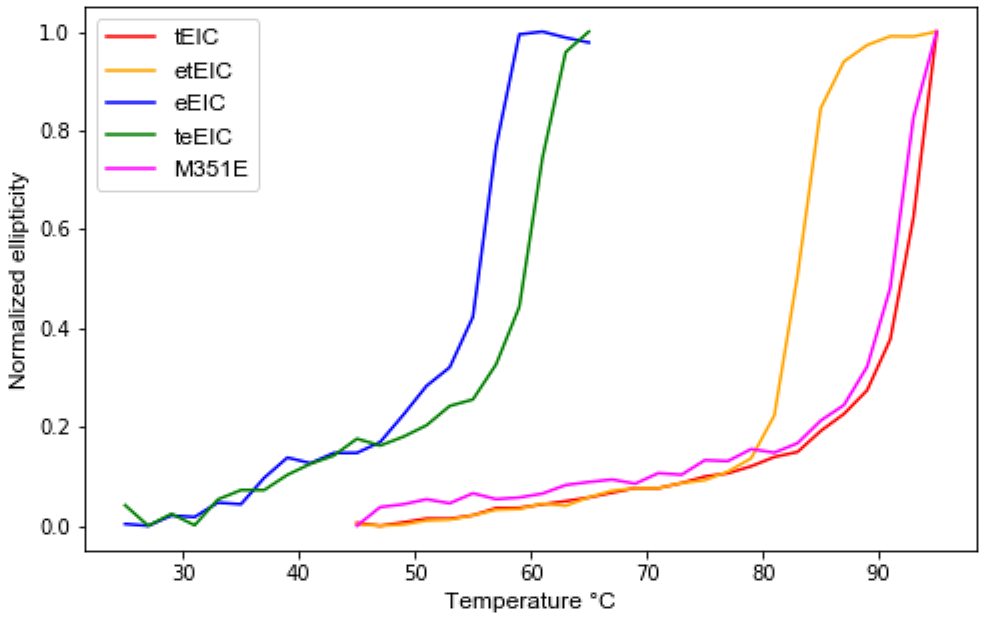


**Fig. S7.** Melting curves for the 4 EIC homologs and M351E. tEIC point Single mutant melting temperatures all fall between etEIC and tEIC. M351E's melting curve is almost identical to wild type tEIC. The M351E mutation has a substantial impact on activity but does not greatly affect stability.

| PC1 | |
|---|---|
| A:ASN:467-A:ASP:464 | 1.000 |
| A:GLU:468-A:LYS:471 | 0.961 |
| A:ARG:465-A:GLN:458 | 0.934 |
| A:ASN:467-A:VAL:470 | 0.852 |
| A:VAL:463-A:VAL:470 | 0.798 |
| A:ARG:465-A:ASN:454 | 0.742 |
| A:GLN:475-A:LEU:461 | 0.737 |

| PC2 | |
|---|---|
| A:LYS:349-A:MET:351 | 1.000 |
| A:GLU:341-A:LYS:349 | 0.924 |
| A:GLN:475-A:TYR:473 | 0.799 |
| A:PHE:295-A:PHE:299 | 0.765 |
| A:ASP:464-B:ASN:352 | 0.751 |
| A:LEU:355-B:ALA:462 | 0.742 |
| A:GLU:350-A:TYR:357 | 0.685 |

| PC3 | |
|---|---|
| A:PHE:354-B:ASN:352 | 1.000 |
| A:TYR:301-A:TYR:344 | 0.884 |

| PC4 | |
|---|---|
| A:TYR:301-A:TYR:344 | 1.000 |
| A:GLY:337-A:LYS:340 | 0.934 |

| | |
|---|---|
| A:ASN:352-B:PRO:353 | 0.782 |
| A:ARG:358-A:ASP:335 | 0.645 |
| A:ARG:304-A:TYR:344 | 0.644 |
| A:GLU:341-A:LYS:349 | 0.614 |
| A:ASN:352-A:LEU:355 | 0.608 |

| | |
|---|---|
| A:ILE:336-A:LEU:334 | 0.925 |
| A:PHE:354-B:ASN:352 | 0.811 |
| A:PRO:343-A:TYR:301 | 0.811 |
| A:PHE:295-A:PHE:299 | 0.786 |
| A:ASN:352-A:GLU:350 | 0.773 |

**Fig. 8.** Top 7 temperature-sensitive contacts on PCs 1-4 sorted according to normalized loading score. Residues highlighted in green are naturally substituted between the homologs.

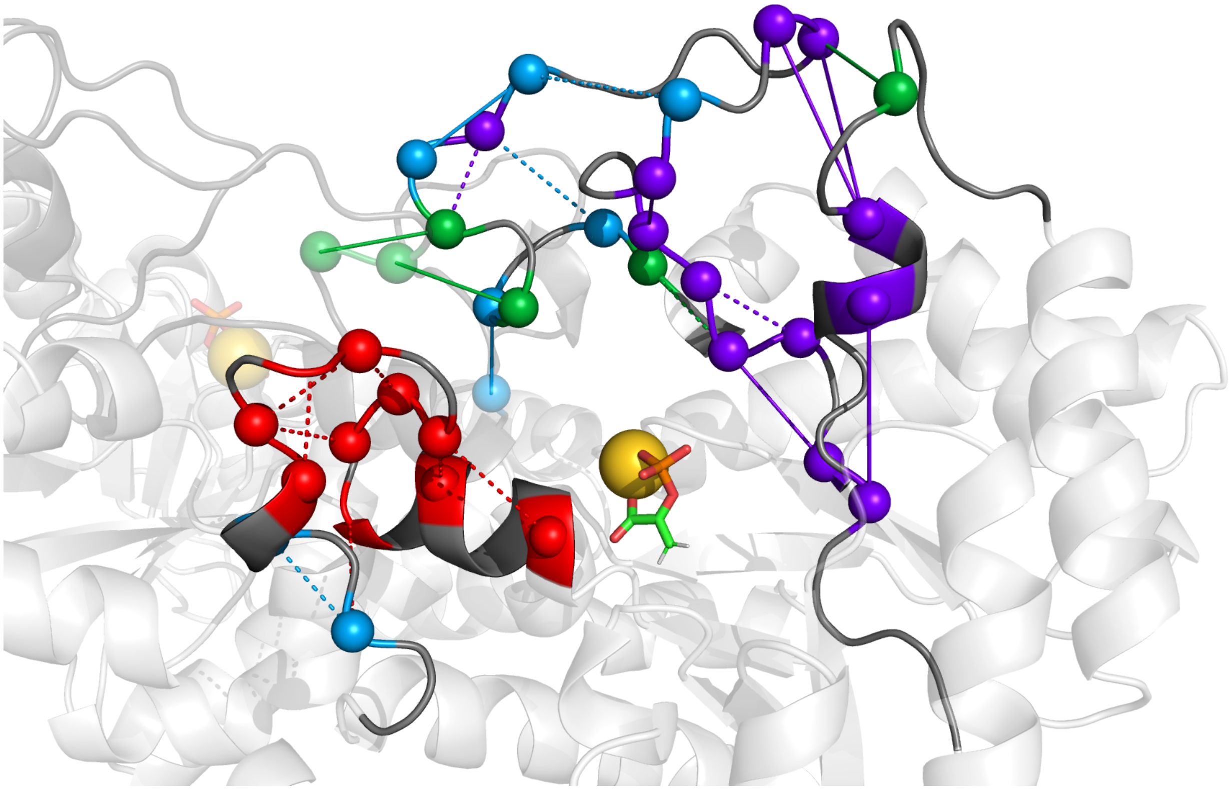

**Fig. S9.** Top temperature sensitive contacts colored according to the PC on which it has the highest loading score. Clear structural clustering is apparent. Contacts that have a different colored sphere on each end appear when a residue has a higher scoring contact with another residue on a different PC. Solid lines indicate contacts with a positive slope of contact frequency across the physiological temperature range (increasing contact frequency) while dashed lines indicate contacts that decrease in contact frequency across the physiological temperature range. PC 3 (green) appears to primarily define the intersubunit interaction between the α3β3 subunits involving conserved residues.

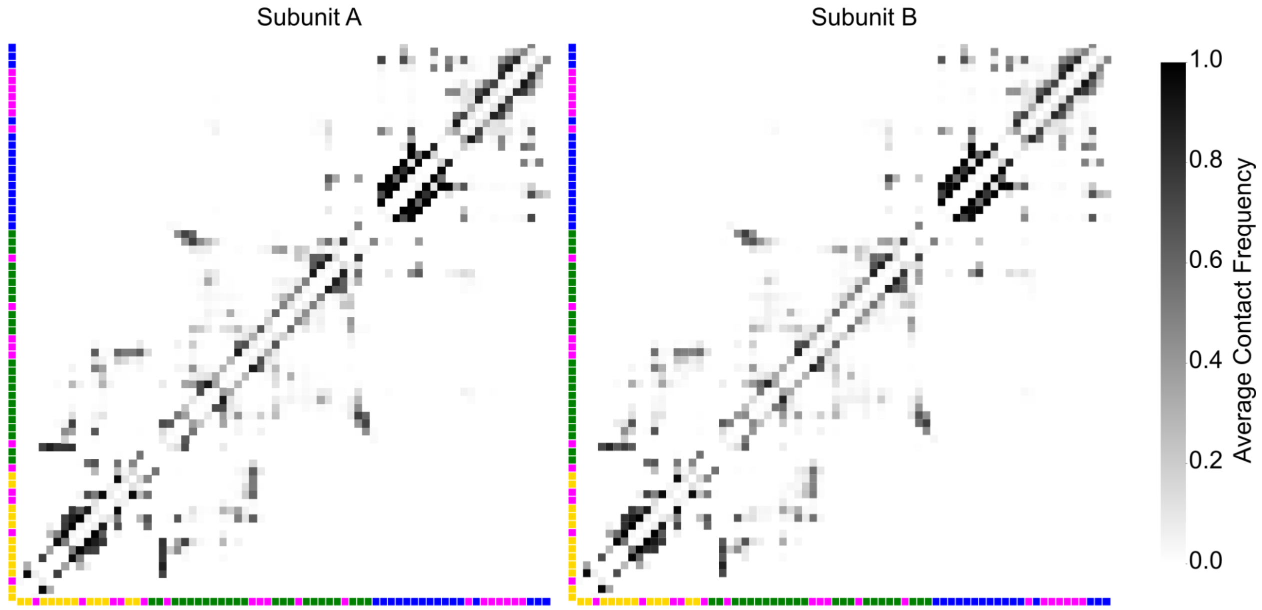


**Fig. S10.** Contact maps for the 2 tEIC subunits A and B depicting contact frequencies obtained from 200ns equilibrium molecular dynamics simulation via Hamiltonain Replica Exchange sampling. An averaged dataset of both subunits (cumulative 400 ns worth of sampling) was used for subsequent analysis.

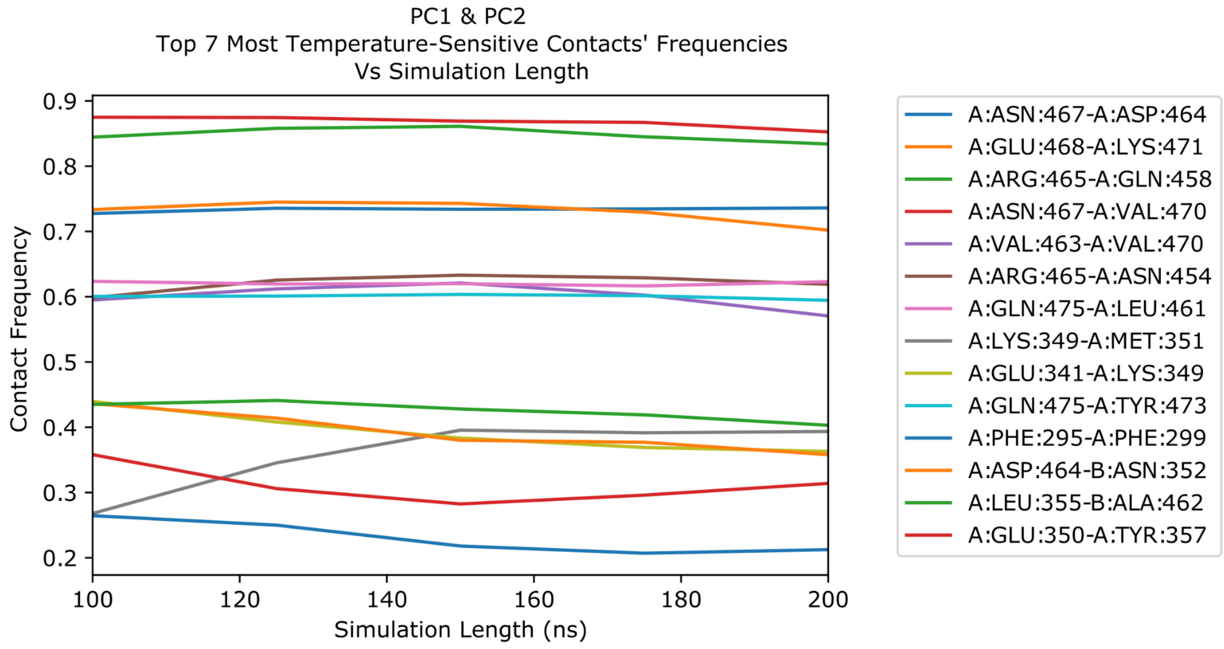


**Fig. S11.** Contact frequencies as a function of simulation length for PC1 and PC2's most temperature-sensitive contacts taken from the simulation system corresponding to a temperature of 62° C (tEIC physiological temperature is 65°C). Stable contact frequencies for these highly sensitive interactions indicates that the simulation is converged for the features of interest.
